# Low-Cost Real-Time Fiber-Optic Distance Sensing for OPM-based Biomagnetic Measurements

**DOI:** 10.64898/2026.10.27.762118

**Authors:** Marco Hübner, Justus Marquetand, Haodi Yang

## Abstract

**Background:** Accurate tracking of the distance between magnetic sensors and biological tissue is particularly relevant to biomagnetic measurements in which stable wearable sensor configurations are not feasible, such as optically pumped magnetometer-based magnetomyography (OPM-MMG), where movement-induced changes in source-to-sensor geometry can nonlinearly affect the recorded signal and complicate motion-artifact control. In OPM setups, such measurements must be performed with strictly nonmagnetic components to avoid noise interference. Therefore, a nonmagnetic, ideally compact approach is needed for real-time monitoring of the sensor-to-source distance.

**Method:** Low-cost, non-magnetic, thin polymer fiber sensors (Keyence FS-N11MN with FU-6F fibers) were evaluated for accurate real-time distance measurement at sub-mm resolution using a fully automated linear stage across a 0–30 mm range and controlled incidence angles between 50° and 90°. The sensors generated analog voltages from 1 V to 5 V, sampled at 2048 Hz, with a 16 ms response time. Custom scripts were used for data analysis, including averaging, noise estimation, and uncertainty weighting.

**Results:** Voltage decreased nonlinearly with distance. For pooled 80°–90° data, weighted Hill1 fits over 0–10 mm and 0–30 mm each yielded an adjusted *R*^2^ greater than 0.9990. Across the five sensors, the mean absolute reconstruction error was 0.0282 mm for 0–10 mm calibration and evaluation and 0.1310 mm when 0–30 mm calibration was evaluated over 10 – 30 mm. A periodic modulation was observed beyond 10 mm, independent of cross-talk, and was consistent with the sensor’s discrete internal signal levels.

**Conclusion:** Low-cost, nonmagnetic polymer fiber sensing may provide a compact, real-time approach for monitoring sensor-to-surface distance over 0–10 mm in OPM-based biomagnetic measurements; extension across the full 0–30 mm range will require range-specific calibration and further validation on skin-facing surfaces.

## 1. Introduction

Biomagnetic measurements can record the weak magnetic fields generated by biological activity and provide non-invasive access to neuronal, cardiac, and muscular electrophysiological processes [1–4]. Since the introduction of superconducting quantum interference devices (SQUIDs) in 1965 by Jim Zimmerman, biomagnetic techniques such as magnetoencephalography (MEG), magnetocardiography (MCG), and magnetomyography (MMG) have become established tools for studying human electrophysiology [2–7]. Historically, those biomagnetic measurements have predominantly relied on SQUID-based systems [3,4]. Although these instruments provide high sensitivity, they require cryogenic cooling to approximately −270 °C using liquid helium [8,9]. Moreover, the SQUID sensing elements are housed in vacuum-insulated dewars [4,10], which create a fixed geometric separation between the sensing elements and the biological tissue. Consequently, the minimal distance between the sensor and the biological tissue (i.e., the sensor-to-source distance) is determined by the physical structure of the dewar and its thermal insulation rather than by flexible sensor placement [3,10,11]. Consistent with the Biot–Savart [12,13] law and prior observations [14,15], even millimeter-scale changes in this distance can alter the measured signal amplitude.

With recent technological advancements, optically pumped magnetometers (OPMs, quantum sensors based on the Zeeman effect) have emerged as a high-sensitivity (10–20 fT 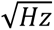) alternative to SQUID systems, enabling flexible sensor placement and wearable measurement configurations [11,16–19]. In contrast to SQUID systems, OPMs operate at room temperature and can be positioned closer to the body surface and adapted to different anatomical regions. This increased flexibility enables new measurement configurations [10,11,20], but it also makes sensor-to-source geometry more variable. In SQUID-based MEG, head position is commonly monitored using head-position coils attached to the participant, whereas comparable tracking is not necessarily integrated into OPM-MEG systems [21–23]. The resulting uncertainty in measurement geometry is particularly relevant to OPM-based MMG because muscle contraction causes the muscle fibers to shorten and bulge, thereby dynamically altering the relative positions of the sensor housing, skin surface, and underlying neuromuscular current sources [15,24–28]. However, in OPM-based MMG measurements, the sensor-to-source distance cannot be directly measured non-invasively during recordings [15]. Tracking the distance between the external OPM housing and the skin can instead provide a proxy for changes in sensor-to-tissue geometry, although it does not measure muscle depth directly. Such monitoring remains technically challenging because components placed near highly sensitive magnetometers must be nonmagnetic. Many conventional inductive, capacitive, and electro-optical displacement probes contain metallic parts or active electronics near the measurement site, which can complicate their integration in magnetically shielded environments [10,11,29–31].

Against this background, a previous study demonstrated the feasibility of fiber-optic distance sensing in OPM-based MMG using a single-sensor approach [14]. That study focused on application feasibility and did not systematically examine sensor-to-sensor variation, angular dependence, calibration over different distance ranges, or output modulation at larger separations. Reflective fiber-optic sensors are suitable for this application because their sensing heads can be nonmagnetic while active electronics remain outside the magnetically sensitive region [29,32]. However, their intensity-based output depends on geometry, surface reflectance, and incidence angle; a calibration obtained in one configuration therefore cannot be assumed to apply unchanged to another [29,32–35]. Accordingly, this study systematically characterizes five low-cost polymer fiber sensors for OPM-compatible distance monitoring by quantifying their voltage–distance response, angular dependence, sensor-to-sensor variability, reproducibility, and calibration performance over a 0–30 mm range. Moreover, the analysis focuses on the near-distance range relevant to OPM-MMG and further examines sensor output behavior at larger separations, including potential systematic modulation.

## 2. Materials and Methods

### 2.1 Sensors and experimental setup

Five available fiber-optic sensing units were used in the present study, each consisting of a Keyence FS-N11MN amplifier and a 4 m polymer optical fiber probe (Keyence FU-6F, Japan; outer diameter 2.2 mm) operated as a reflective distance sensor. The metallic ferrule at the fiber end was removed to ensure nonmagnetic operation. The sensors emitted red light at 630 nm and provided a 1–5 V analog output, which was recorded with a signal acquisition system (OT Bioelettronica Quattrocento, Italy). All five sensors were supplied in parallel with 12 V DC (maximum current 1.5 A). MEGA is the manufacturer-defined 16 ms response-time setting of the FS-N11MN amplifier and was used for all measurements; faster response-time settings were not evaluated in this study. The analog scaling was set to 8500 mV. A CNC linear stage (SBR16 with SFU1605 ball screw, 5 mm pitch; RATTMOTOR, China) driven by a NEMA 23 stepper motor was used to position the target surface (Figure 1A). The target consisted of a flat polylactic acid (PLA) plate covered with 0.3 mm white paper (Figure 1A).

**Figure 1:**
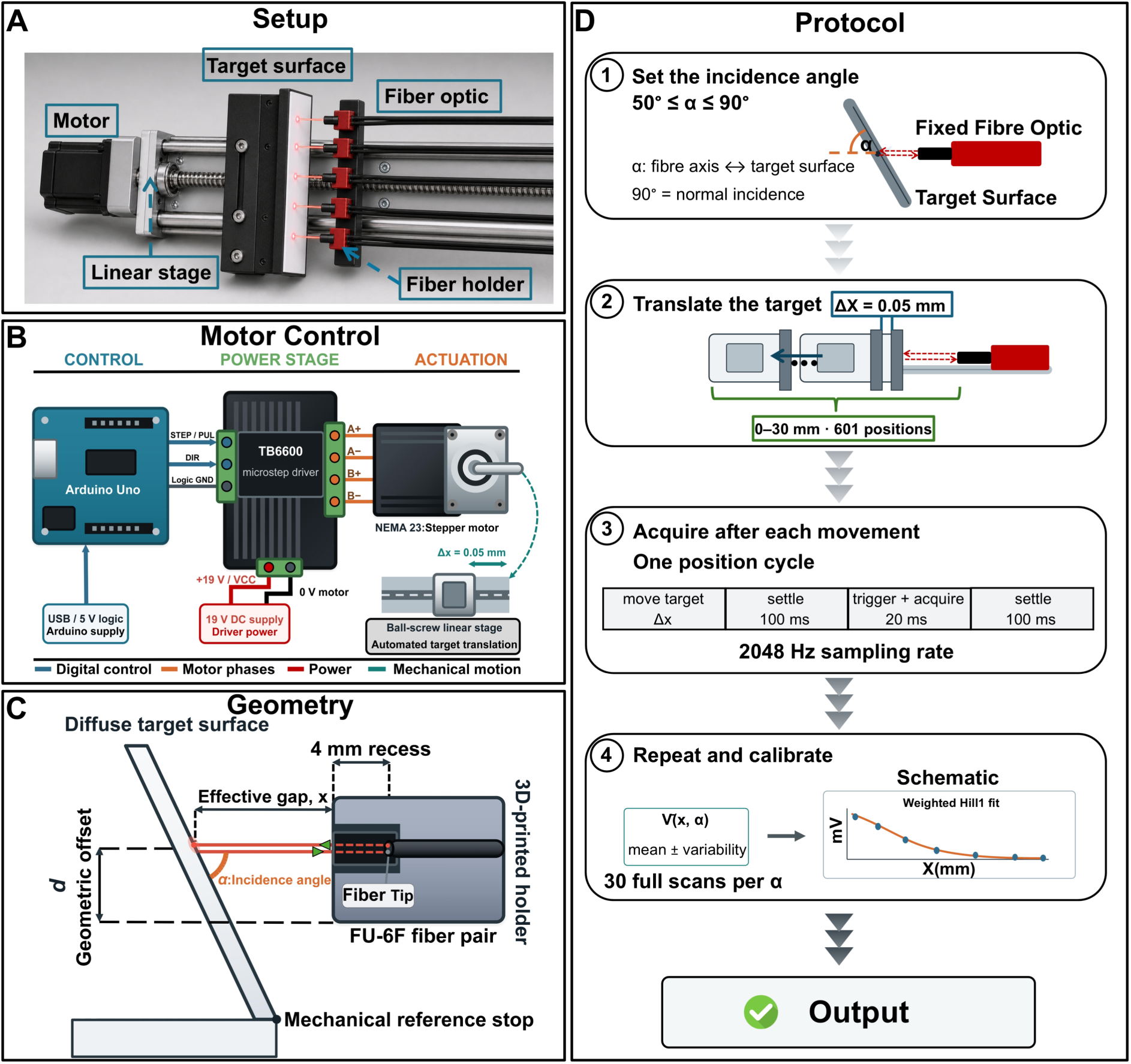
Experimental setup and automated protocol for angle-resolved calibration of the fiber-optic distance-sensing system. **(A)** Physical setup with five fixed FU-6F polymer fiber probes mounted in a custom black polylactic acid holder and a diffuse target translated by a ball-screw linear stage. **(B)** Motor-control architecture: an Arduino Uno supplied step and direction signals to a TB6600 microstep driver controlling a National Electrical Manufacturers Association 23 stepper motor. **(C)** Fiber–target geometry. Each fiber tip was recessed by 4 mm; the incidence angle α was defined between the fiber axis and target surface, with α = 90° corresponding to normal incidence. The geometric offset d between the fiber center and mechanical reference stop was used to calculate the effective gap x. **(D)** Automated protocol. At each incidence angle from 50° to 90°, the target was translated from 0 to 30 mm in 0.05 mm steps (601 positions). Each movement was followed by 100 ms settling, 20 ms triggering and acquisition, and a further 100 ms settling period. Signals were sampled at 2048 Hz, and 30 full scans were acquired per angle before averaging and weighted Hill1 fitting.

### 2.2 Experimental protocol and geometric configuration

The motor was controlled by an Arduino Uno microcontroller (Arduino Uno, Italy) and a TB6600 stepper-motor driver (OUYZGIA stepper motor driver TB6600, China). After each movement, the system paused for 100 ms, sent a 5 V digital trigger pulse of 20 ms to the acquisition system, and a further 100 ms pause preceded the next movement. Each full scan covered 601 positions over 0–30 mm in 0.05 mm steps and was repeated 30 times at each incidence angle. The automated sequence is summarized in Figure 1D. A 2-min cooling pause was inserted after every five scans to limit heating of the motor and driver.

To assess whether simultaneous sensor operation introduced optical or electronic cross-talk, the measurement at 90° was repeated over 0–30 mm with only Sensor 1 active and Sensors 2–5 disconnected. This isolated-sensor measurement followed the same automated protocol as the main experiment, including 0.05 mm steps, 601 measurement positions, and 30 repeated scans, under otherwise unchanged geometric and illumination conditions. The fibers were mounted in a custom black polylactic acid holder printed with an UltiMaker S7 (Netherlands). The holder was designed to minimize parasitic reflections, defined here as light returned from the holder walls or nearby structures rather than from the target. Each fiber tip was recessed by 4 mm to place the intrinsic near-field blind zone within the holder and obtain a monotonic voltage–distance relation over the external measurement range (Figure 1C). Signals were recorded with OT Biolab+ at 2048 Hz, exported in MATLAB format, processed with a custom Python script, and fitted in OriginLab 2025b (OriginLab, USA).

Measurements were performed at incidence angles from 50° to 90° to quantify how departures from normal incidence affected the voltage–distance relation. The incidence angle α was defined between the fiber axis and target surface, with α = 90° corresponding to normal incidence (Figure 1C). Angles below 50° were not investigated because angle dependence was already evident at 50°. For Sensor 1, linear interpolation showed that 2500 mV corresponded to 4.52 mm at 90° and 3.28 mm at 50°, a difference of 1.24 mm. Because this difference exceeded the targeted submillimeter precision in the near range, the 50° dataset was excluded from pooled calibration and the main calibration figures. Nominal stage displacement and corrected effective distance were treated separately. The effective distance *x* was the geometrically corrected separation between the holder and illuminated target, including the offset *d* between the fiber center and mechanical reference stop (Figure 1C).

### 2.3 Sensor configuration and standardization

For each FS-N11MN unit, the light emission was set manually while the target was oriented at 90° and at a nominal separation of 0 mm. The goal was a received intensity of about 7300 arbitrary units. This target value was selected based on preliminary testing, as the manufacturer does not specify an optimal calibration intensity for the FS-N11MN amplifier. In MEGA mode, the maximum displayable received intensity is 9999 arbitrary units (a.u.) under standard display settings. A level of approximately 7300 a.u. was chosen to avoid operation close to the upper display limit. The emission was adjusted accordingly and Table 1 lists the emission percentage together with the corresponding readback intensity for each sensor–fiber pair. Each unit was permanently paired with its specific two-core polymer fiber, and the emitting and receiving cores were fixed by labeling to avoid accidental swaps that would change the absolute signal level. Swapping receiving cores was observed to alter the absolute signal level. Although this effect was not quantified systematically, permanent pairing and labeling were implemented to ensure consistent measurement conditions.

**Table 1:** Sensor-specific emission settings and corresponding received-intensity readbacks at normal incidence. For each sensor–fiber pair, the emission level was manually adjusted at an incidence angle of 90° and a nominal distance of 0 mm to obtain a received intensity of approximately 7300 arbitrary units (a.u.). The selected emission setting was subsequently kept unchanged throughout all measurements. Each amplifier remained permanently paired with its designated fiber probe.

| Sensor ID | Emission setting [%] | Received intensity [a.u.] |
| --- | --- | --- |
| 1 | 17 | 7441 |
| 2 | 32 | 7329 |
| 3 | 13 | 7214 |
| 4 | 4 | 7281 |
| 5 | 9 | 7232 |

The emission setting for each FS-N11MN unit was defined once at the beginning of the experiments and was not readjusted on subsequent measurement days. The fiber ends remained fixed throughout the study. For different incidence angles, only the target surface was repositioned to maintain the defined mechanical reference point. No separate reference measurement was performed to quantify potential changes in absolute received intensity between sessions. All calibration fits were performed directly on the recorded voltage–distance data.

### 2.4 Distance correction, signal processing and calibration

The effective distance was calculated as:

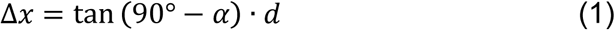

where is the angle-dependent distance correction in millimeters, is the incidence angle between the fiber axis and the target surface, and is the fixed geometric offset between the fiber center and the mechanical reference stop. The correction was added to the nominal stage displacement to obtain the effective fiber-to-target distance.

For each sensor and position, voltage samples were grouped into trigger events, yielding 40 data points per window. Each measurement cycle covered 601 positions over 0–30 mm in 0.05 mm steps. The mean voltage within each trigger represented the signal at the corresponding distance. Intra-trigger noise was quantified as the standard deviation of samples within each trigger window and was averaged across the 30 repeated scans at each distance. Within-protocol repeatability was assessed as the standard deviation of the mean voltages across the 30 repeated full scans. The standard error of the mean for each trigger event was calculated as:

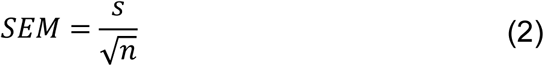

where *M* is the standard error of the mean, is the standard deviation of the samples and the number of data points per trigger window. In addition, for each distance the mean voltage across all cycles and the corresponding standard deviation were determined, allowing a direct comparison of noise within single triggers and variability across repeated measurements.

The relationship between monitor-output voltage and effective distance was modeled using a four-parameter Hill1 function:

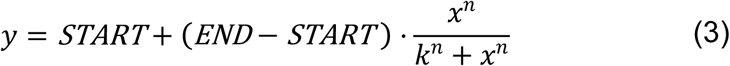

where is the fitted monitor-output voltage in millivolts, is the effective fiber-to-target distance in millimeters, *START* is the fitted voltage at = 0, and *END* is the asymptotic voltage approached at larger distances. The parameter is the half-response distance, at which the fitted voltage lies halfway between *START* and *END*, and is the Hill coefficient controlling the steepness and shape of the fitted curve. The model was fitted by weighted least squares using the inverse variance derived from the combined measurement uncertainty. Instrumental weighting was applied using the combined uncertainty as inverse variance weighting factor. Fits were performed for three angular ranges (80–90°, 70–90°, and 60–90°) and for two distance intervals: 0–10 mm and 0–30 mm. The 50° dataset was excluded from the calibration fits because it showed pronounced systematic deviations. For example, at a voltage level of 2500 mV the inferred distance differed by approximately 1.24 mm compared to the 90° reference, exceeding the targeted submillimeter precision in the near range. The inverse calibration function was used for voltage-to-distance back-calculation, and voltage values outside the valid range of the fitted inverse function were documented as definition gaps.

### 2.5 Statistical analysis

Statistical analyses were conducted in Python (version 3.12.14). Friedman tests and Wilcoxon signed-rank tests were performed using SciPy (version 1.18.1), whereas pandas (version 3.0.6) and NumPy (version 2.5.3) were used for data organization and numerical computation, respectively. For each sensor, reconstruction error was defined as reconstructed distance minus nominal distance. Mean absolute error (MAE), root mean squared error (RMSE), mean bias, and maximum absolute error were calculated for each calibration and evaluation condition. Descriptive results are reported as the mean, standard deviation, median, minimum, and maximum across the five sensors.

Sensor-related differences were evaluated using block-level mean absolute error as the primary outcome. For each held-out scan index from scans 26–30, errors from the 80° and 90° datasets were pooled, resulting in five paired technical blocks per sensor and condition. The five sensors were compared using the nonparametric Friedman test for paired observations. Because only five paired blocks were available, exact randomization *p* values were used for primary inference. Kendall’s *W* was reported as an effect-size measure. When the overall Friedman test was significant, two-sided paired Wilcoxon signed-rank tests were used for pairwise sensor comparisons. Values of *p* from the ten pairwise comparisons within each condition were adjusted using the Holm method. Statistical significance was defined as an adjusted two-sided *p* value below 0.05.

## 3. Results

### 3.1 Angle-dependent response and Hill1 calibration

Across the five sensor–fiber pairs, mean output decreased nonlinearly across the 0–30 mm sweep, with a smaller distance-dependent modulation superimposed at larger separations. Figures 2 and 3 show Sensor 1; corresponding results for Sensors 2–5 are provided in the Supplementary Figure 1–4 and show a broadly comparable qualitative response with sensor-specific absolute output levels.

**Figure 2:**
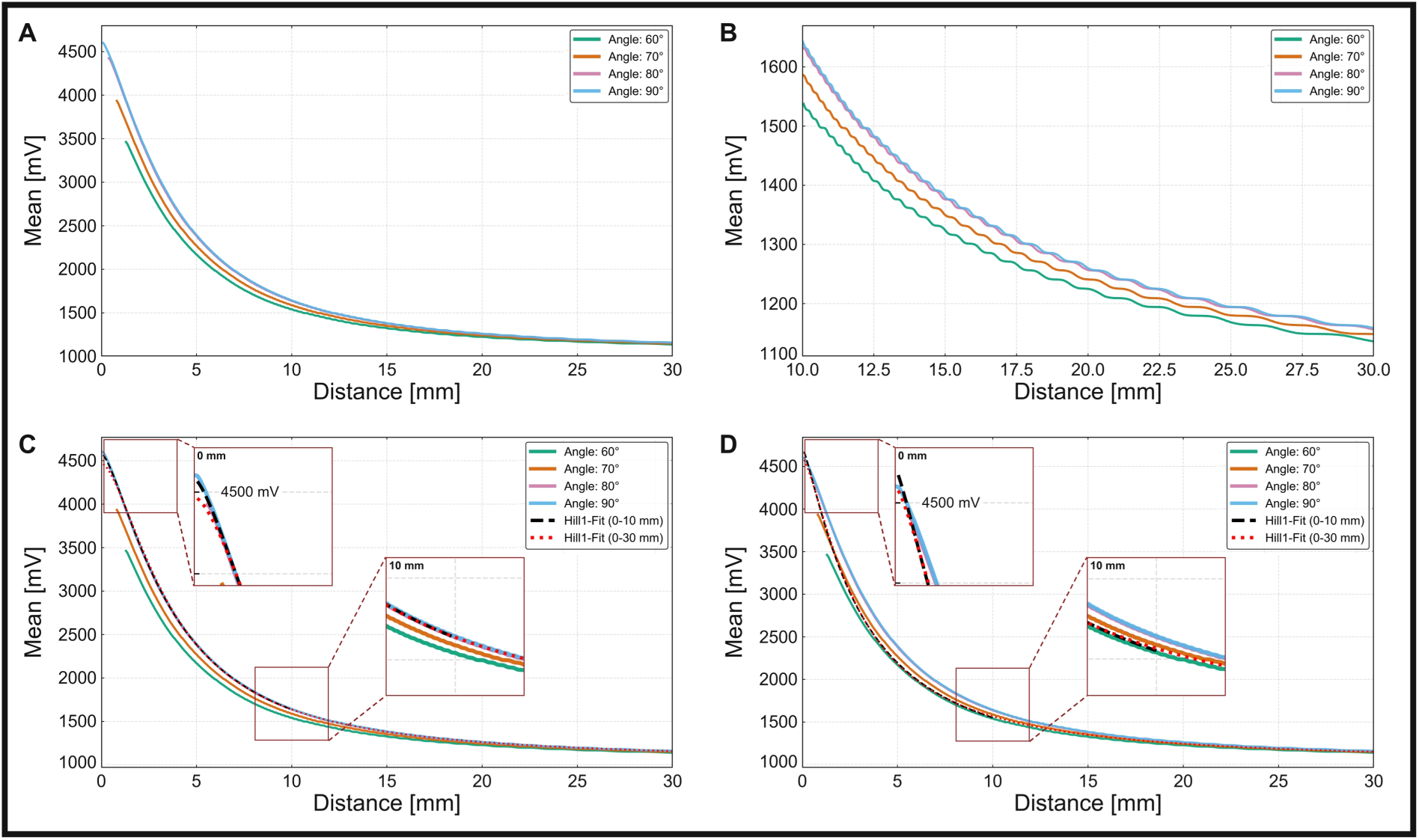
Angle-dependent voltage response and Hill1 calibration for Sensor 1. The figure shows the mean monitor-output voltage measured over a distance range of 0–30 millimetres (mm) at incidence angles of 60°, 70°, 80° and 90°, together with the corresponding Hill1 calibration curves. **(A)** Mean voltage–distance responses over 0–30 mm. **(B)** Enlarged view of the 10–30 mm range, showing the distance-dependent modulation at larger separations. **(C)** Mean responses with uncertainty-weighted Hill1 fits derived from the pooled 80° and 90° data. **(D)** Mean responses with uncertainty-weighted Hill1 fits derived from the pooled 60°–90° data. Black dashed lines show fits over 0–10 mm, whereas red dotted lines show fits over 0–30 mm. In panels C and D, the dark-red boxes mark the regions enlarged in the insets, and the dashed connector lines link each selected region to its corresponding inset. The upper insets enlarge the high-voltage response around the nominal 0 mm position, whereas the lower insets enlarge the region around 10 mm to show the relationship between the measured and fitted curves at the boundary of the near-range calibration interval. The measured 50° data are not shown because, for Sensor 1, a monitor output of 2500 mV corresponded to a distance 1.24 mm shorter than at 90°; this dataset was therefore excluded from the pooled fits.

**Figure 3:**
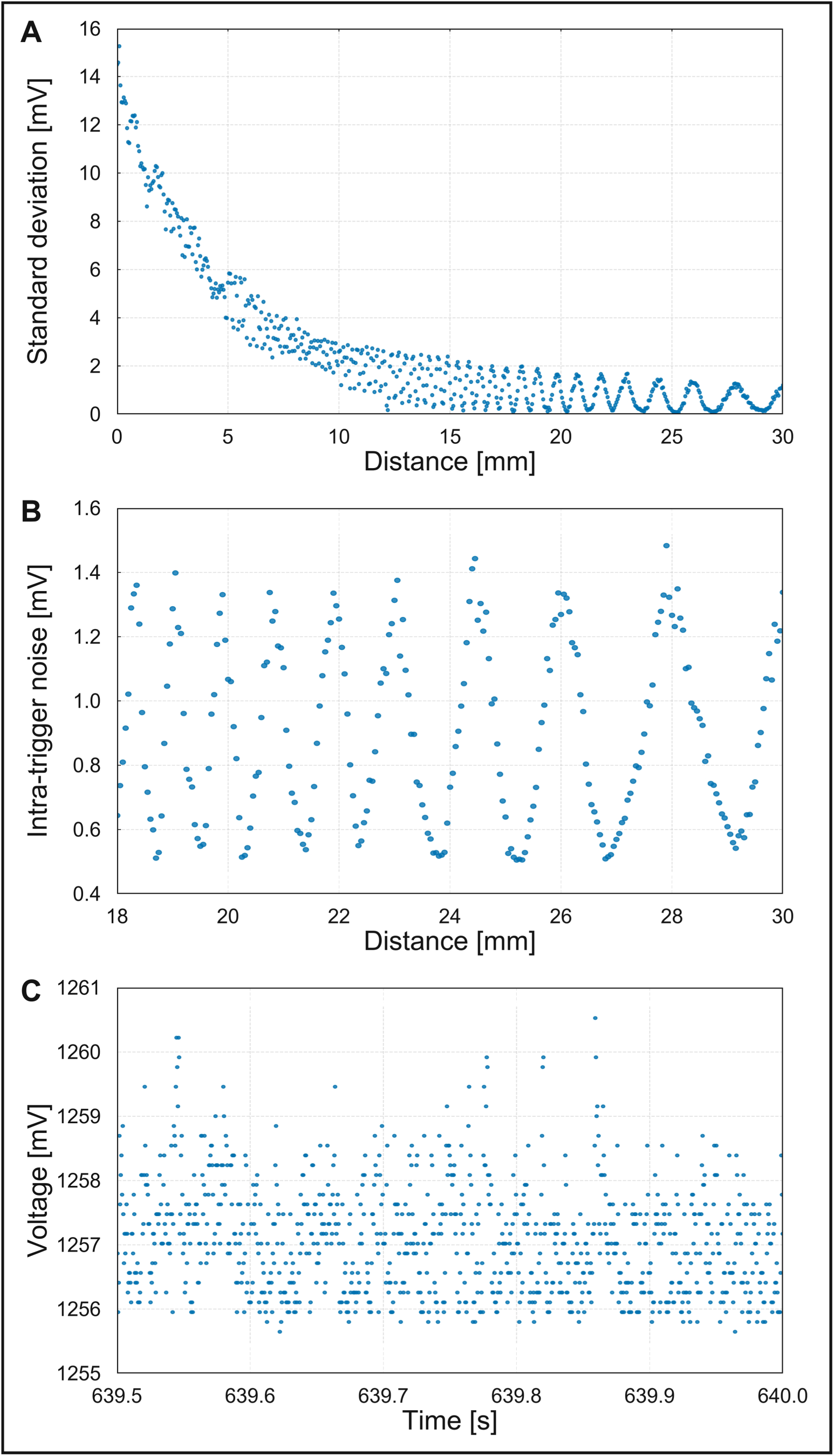
Variability and time-resolved output of Sensor 1 at 90°. **(A)** Standard deviation of the mean voltage across 30 repeated scans at each distance over 0–30 mm. **(B)** Intra-trigger noise, calculated as the standard deviation of approximately 40 samples within each 20 ms trigger window, shown for 18–30 mm. **(C)** A 0.5 s segment of the monitor-output voltage at 20 mm, showing recurrent discrete voltage levels. All panels were derived from the isolated-sensor measurement in which only Sensor 1 was active.

Figure 2A shows the mean Sensor 1 output at incidence angles of 60°, 70°, 80°, and 90°. Because the effective distance incorporated an angle-dependent geometric correction for the offset between the fiber center and the mechanical reference stop, the first sampled effective distance differed among angles (60°: 1.30 mm; 70°: 0.82 mm; 80°: 0.40 mm; and 90°: 0.00 mm). At these initial positions, the mean voltage ranged from 3461.85 to 4603.65 mV; near 30.00 mm, it ranged from 1136.07 to 1158.65 mV. Thus, the voltage decreased by approximately 2325.78–3445.00 mV across the measured range, depending on the incidence angle. The decrease was steepest near the target and progressively flattened with increasing distance, consistent with a sigmoidal Hill-type response. The separation between the angle-dependent curves was greatest in the near range and narrowed toward 30 mm. In the enlarged 10–30 mm view, the superimposed modulation became more apparent, and the spacing between successive modulation features increased with distance (Figure 2B).

Weighted Hill1 fits were calculated from the pooled 80°–90° data (Figure 2C) and the pooled 60°–90° data (Figure 2D) over 0–10 and 0–30 mm. For the pooled 80°–90° data, the adjusted *R*² values were 0.9998 for the 0–10 mm fit and 0.9997 for the 0–30 mm fit. However, the inverse functions left definition gaps of 0.20 and 0.45 mm, respectively, at 90°. For the pooled 60°–90° data, the corresponding adjusted *R*² values were 0.9602 and 0.9770, and no definition gaps occurred. The enlarged regions around the nominal 0 and 10 mm positions further illustrate these differences. In Figure 2C, the fitted curves closely followed the near-perpendicular 80° and 90° responses, although the highest voltage values near the nominal 0 mm position extended beyond the valid range of the inverse calibration. In Figure 2D, the fitted curves represented a compromise across the wider 60°–90° angular range and covered the measured voltage range without definition gaps. Thus, the near-perpendicular calibration provided a closer fit to the 80°–90° data, whereas the wider-angle calibration provided broader angular coverage at the expense of lower fit quality.

### 3.2 Far-range signal modulation and variability

In the isolated-sensor measurement at 90°, performed with Sensors 2–5 disconnected, the far-range modulation remained visible in the mean voltage, between-scan variability, and intra-trigger noise. This observation suggests that cross-talk between the five sensor channels was unlikely to be the principal cause of the modulation. The standard deviation across 30 scans was 14.50 mV at 0 mm and declined to no more than 2.17 mV beyond

15 mm, where the distance-dependent modulation became apparent (Figure 3A). Between 18 and 30 mm, intra-trigger noise ranged from 0.51 to 1.48 mV and showed a corresponding periodic pattern (Figure 3B). At 20 mm, a 0.5 s segment of the Sensor 1 output occupied recurrent voltage levels between 1255.65 and 1260.53 mV rather than a single constant level (Figure 3C).

Corresponding mean curves, calibration fits, variability analyses, and time-domain plots for Sensors 2–5 are provided in the Supplementary Figure 1–4. These sensors showed comparable nonlinear distance dependence and far-range modulation, although their absolute output levels and reconstruction errors differed.

### 3.3 Independent distance reconstruction across sensors

Distance reconstruction was evaluated for five sensors using scans 26–30, which were excluded from calibration. For Figure 4, mean absolute error, root mean squared error, and maximum absolute error were each divided by the largest value of that metric across the 20 sensor–condition combinations. Mean bias was divided by the largest absolute bias while retaining its sign. This normalization was used only for visualization; statistical analyses used the original values in millimeters (Supplementary Table 1).

**Figure 4:**
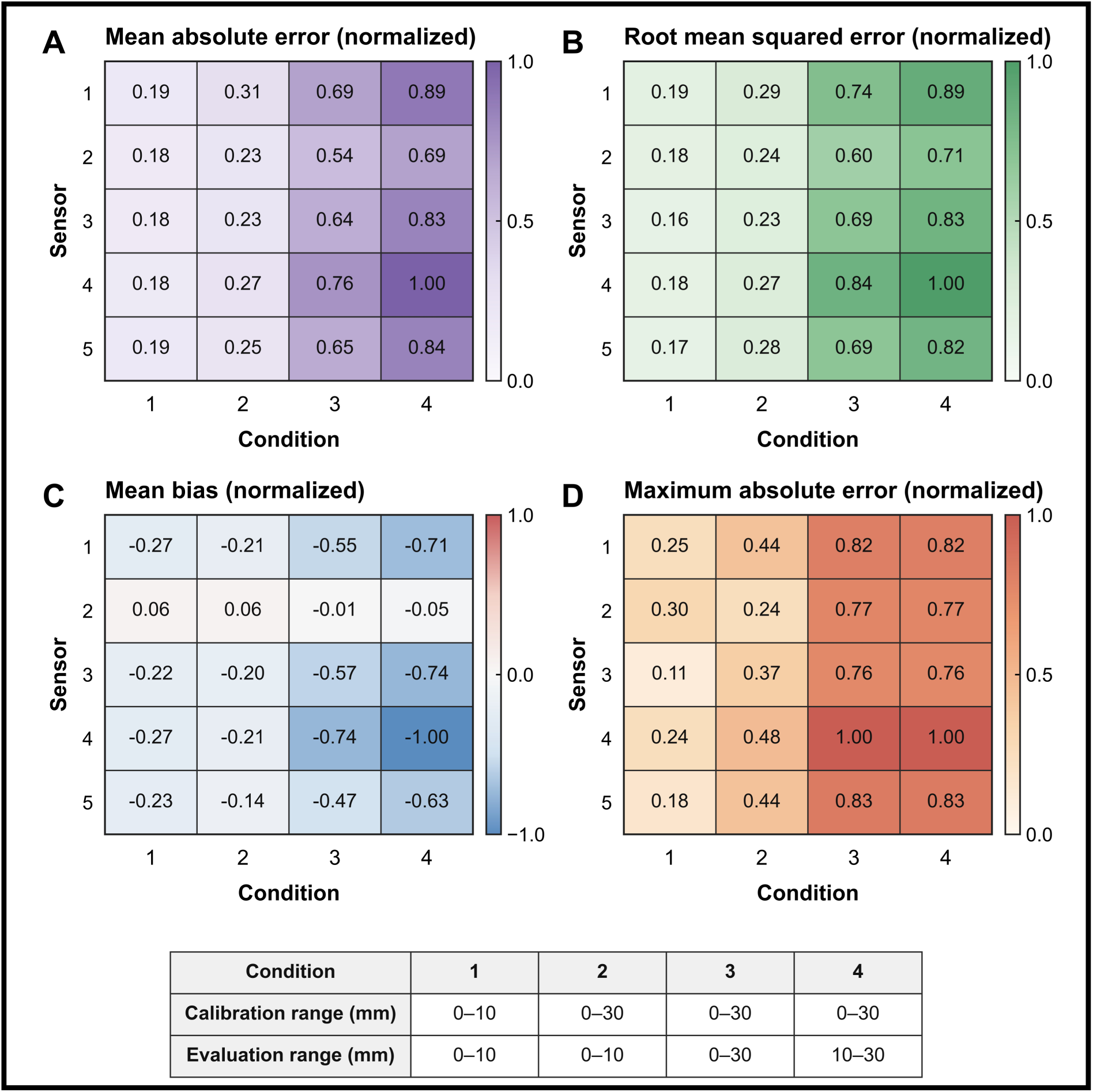
Normalized distance-reconstruction errors across five sensors under different calibration and evaluation ranges. Distance reconstruction was assessed using scans 26–30 as independent test data after calibration with scans 1–25 from the 80° and 90° measurements. **(A)** Normalized mean absolute error, representing the average magnitude of the reconstruction errors. **(B)** Normalized root mean squared error, which gives greater weight to larger reconstruction errors. **(C)** Normalized mean bias, defined as the reconstructed distance minus the nominal distance; negative values indicate an overall tendency to underestimate distance. **(D)** Normalized maximum absolute error, representing the largest reconstruction error observed for each sensor and condition. Conditions 1–4 correspond to calibration/evaluation ranges of 0–10/0–10, 0–30/0–10, 0–30/0–30 and 0–30/10–30 millimeters, respectively. For panels A, B, and D, each value was divided by the maximum value of that metric across all sensors and conditions. For panel C, each bias was divided by the largest absolute bias and its sign was retained. The cell labels and color bars therefore show unitless normalized values; the original values in millimeters are reported in Supplementary Table 1.

The normalized heat maps emphasize the same range-dependent pattern as the unscaled errors in Figure 4 and Supplementary Table 1. Across the five sensors, mean absolute error increased from 0.0282 mm under Condition 1 (0–10 mm calibration and evaluation) to 0.1310 mm under Condition 4 (0–30 mm calibration evaluated over 10–30 mm). Over the same conditions, root mean squared error increased from 0.0358 to 0.1724 mm and maximum absolute error from 0.1656 to 0.6426 mm. Mean bias was −0.0182, −0.0140, −0.0460, and −0.0616 mm under Conditions 1–4, respectively, indicating greater underestimation over the wider evaluation ranges. The Friedman test found no evidence of an overall sensor effect under Condition 1 (χ² (4) = 3.04, exact *p* = 0.594, Kendall’s *W* = 0.152), but detected an overall sensor effect under Conditions 2–4 (all exact *p* < 3 × 10⁻ ⁷; *W* = 0.952–0.968). No individual sensor pair remained significant after Holm adjustment (all adjusted *p* ≥ 0.625).

## 4. Discussion

This study examined whether a cost-effective, nonmagnetic, reflective fiber sensor can serve as a calibrated distance proxy for OPM-based MMG recordings. Our findings suggest that the system may provide a compact, real-time distance proxy for 0–10 mm, while extending it across the full 0–30 mm range would require range-specific calibration and further validation on skin-facing surfaces. However, based on previous studies regarding the attenuation of the biomagnetic signal with distance [14,15], we consider a range of 0–10 mm to be optimal. Across the five sensor–fiber pairs, the near-range response was reproducible and well described by Hill1 fits, whereas angular dependence and periodic far-range modulation constrained calibration outside the intended operating conditions.

### 4.1 Calibration characteristics and operating considerations

The five FS-N11MN sensor–fiber pairs required emission settings between 4% and 32% to reach a similar received intensity at 0 mm and 90°. This spread indicates that absolute scaling cannot be transferred reliably between nominally identical pairs. Differences in fiber routing, end-face preparation, coupling, or holder alignment could contribute, but these factors were not varied independently. The present data therefore support permanent pairing of each amplifier and fiber and an individual calibration for each pair.

The 4 mm recess moved the intrinsic near-field blind zone into the holder geometry and produced a monotonic external response on the diffuse target. Within 80°–90°, both Hill1 fits closely represented the measured response, whereas pooling 60°–90° reduced fit quality but avoided the short definition gaps. This trade-off, highlighted by the enlarged regions around 0 and 10 mm in Figure 2C and 2D, indicates that the calibration angle range should be selected according to the expected variation in sensor–surface alignment rather than on fit quality alone. The operating-angle range should therefore be specified before calibration: a near-perpendicular calibration is more representative when alignment is constrained, while a wider-angle calibration may be preferable when the incidence angle varies. Because a single intensity value does not identify distance and angle independently, smaller angles require separate calibration rather than extrapolation from the near-perpendicular fit.

The far-range modulation was observed in the mean response, between-scan variability, and intra-trigger noise and persisted when Sensor 1 was operated alone. Cross-talk between sensors is therefore unlikely to explain the pattern. The recurrent voltage levels in the time-domain trace are compatible with output discretization or internal amplifier processing, while multimode-fiber coupling could also contribute [36]. These possibilities remain hypotheses because the amplifier processing is proprietary and the mechanisms were not tested separately.

The calibrated near range overlaps the millimetre-scale housing-to-skin changes reported during OPM-MMG recordings [14], but the present measurements used a homogeneous white diffuse target rather than skin. Reflectance, pigmentation, perfusion, sweat, hair, and surface texture may alter the received intensity [37–40], so validation on skin-facing surfaces is needed before application during human recordings. In addition, no independent reference measurement was collected across measurement days; between-session stability of absolute intensity was therefore not quantified. MEGA denotes the 16 ms response-time setting used here. Faster amplifier settings were not compared, and the present results should not be extrapolated to high-frequency tissue motion. Together with pair-specific scaling and unresolved angle–distance ambiguity, these factors define the current operating limits.

### 4.2 Distance-dependent reconstruction and inter-sensor variability

Across the five sensors, reconstruction errors were smaller within 0–10 mm than over the wider evaluation ranges (Figure 4 and Supplementary Table 1). This pattern is consistent with the voltage–distance curve: its steeper near range provides a larger voltage change per unit displacement, whereas the flatter distal range converts a given voltage fluctuation into a larger distance error. The difference between Conditions 1 and 2 also indicates that the selected fitting range can influence near-range reconstruction. A 0–10 mm calibration is optimized for the most sensitive part of the response, whereas a 0–30 mm calibration must accommodate both the steep proximal and flatter distal regions. Range-specific models, weighted fitting, or separate functions for different distance intervals may therefore improve future calibration procedures.

The increasingly negative mean bias over wider evaluation ranges indicates systematic underestimation rather than only greater random error. Sensor 2 had a bias closer to zero, which may reflect closer agreement between its fitted curve and held-out measurements or a more even balance of positive and negative errors. Because signed errors can cancel, bias should be interpreted together with mean absolute error and root mean squared error. The overall Friedman tests detected sensor effects under Conditions 2–4, but no pairwise comparison remained significant after Holm adjustment. Sensor-specific conclusions should therefore remain cautious, particularly because inference was based on five paired technical blocks. Small differences in fiber positioning, optical coupling, detector response, or fitted parameters could contribute to the observed variation. The results are consistent with individual calibration of each sensor and with prioritizing the near range for applications that monitor small sensor-to-surface separations.

## 5. Conclusion

A nonmagnetic fiber-optic distance-sensing configuration was characterized for use in magnetically sensitive environments. Reconstruction errors were lowest over 0–10 mm, whereas distance-dependent modulation and higher errors were observed over 10–30 mm. These findings support polymer fiber sensing as a compact, low-cost method for near-range distance monitoring in wearable optically pumped magnetometer systems, subject to sensor-specific calibration and validation on skin-facing surfaces.

## 6. Declarations

### 6.1 Data availability statement

The original data can be made accessible upon reasonable request directed to the corresponding author.

### 6.2 Ethics statement

This study did not involve human participants, animals, or identifiable personal data; therefore, ethics approval and informed consent were not required.

### 6.3 Author contributions

MH, JM and HY: study design, data collection and analysis, writing, and revision. All authors contributed to the article and approved the submitted version.

## Abbreviations

OPM: Optically pumped magnetometer
MEG: Magnetoencephalography
BITS: Center for Bionic Intelligence Tübingen Stuttgart
MMG: Magnetomyography
SQUID: Superconducting quantum interference device
MCG: Magnetocardiography
EMG: Electromyography
OPM-MMG: Optically pumped magnetometer-based magnetomyography
DC: Direct current
CNC: Computer numerical control
NEMA: National Electrical Manufacturers Association
PLA: Polylactic acid
a.u.: Arbitrary units
ID: Identification
MAE: Mean absolute error
RMSE: Root mean squared error

