## Supplemental Materials for "Low-Cost Real-Time Fiber-Optic Distance Sensing for OPM-based Biomagnetic Measurements"

Supplementary Figure 1 — Sensor 2

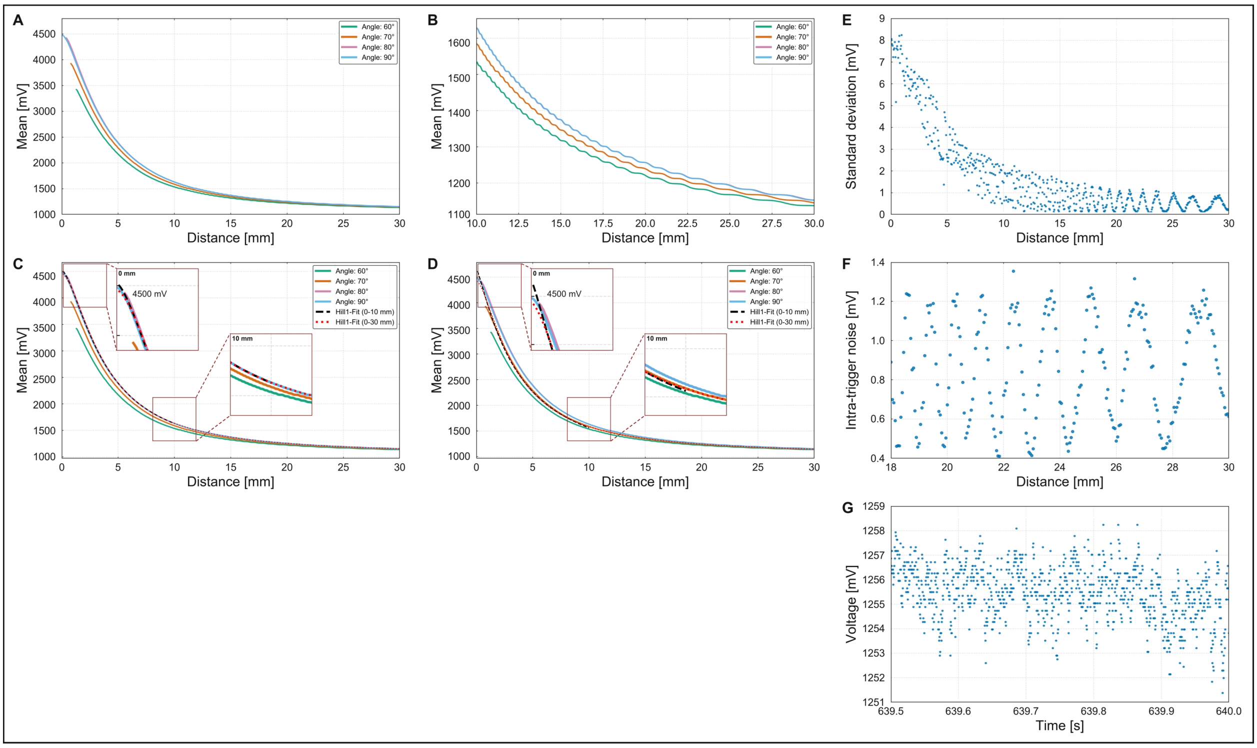


***Supplementary Figure 1:*** Angle-dependent voltage response, Hill1 calibration, and signal variability for Sensor 2. Voltage, distance, and time are reported in millivolts (mV), millimetres (mm), and seconds (s), respectively. **(A)** Mean monitor-output voltage across 30 repeated scans over 0–30 mm at incidence angles of 60°, 70°, 80°, and 90°. **(B)** Enlarged view of the 10–30 mm range, showing the distance-dependent modulation at larger separations. **(C)** Mean voltage–distance responses with uncertainty-weighted Hill1 fits derived from the pooled 80° and 90° data. **(D)** Mean responses with uncertainty-weighted Hill1 fits derived from the pooled 60°–90° data. Black dashed lines represent fits over 0–10 mm, whereas red dotted lines represent fits over 0–30 mm. In panels C and D, the dark-red boxes mark the selected regions, the dashed connectors link these regions to the corresponding insets, and the upper and lower insets enlarge the regions around the nominal 0 and 10 mm positions, respectively. **(E)** Standard deviation of the mean voltage across 30 repeated scans at each distance over 0–30 mm at 90°. **(F)** Intra-trigger noise, calculated as the standard deviation of approximately 40 samples within each 20 ms trigger window and averaged across the 30 scans, shown over 18–30 mm at 90°. **(G)** A 0.5 s segment of the monitor-output voltage at 20 mm and 90°, showing the sampled voltage levels over time. The measured 50° data are not displayed because these measurements were excluded from the pooled calibration fits according to the criterion described in the Methods.

Supplementary Figure 2 — Sensor 3

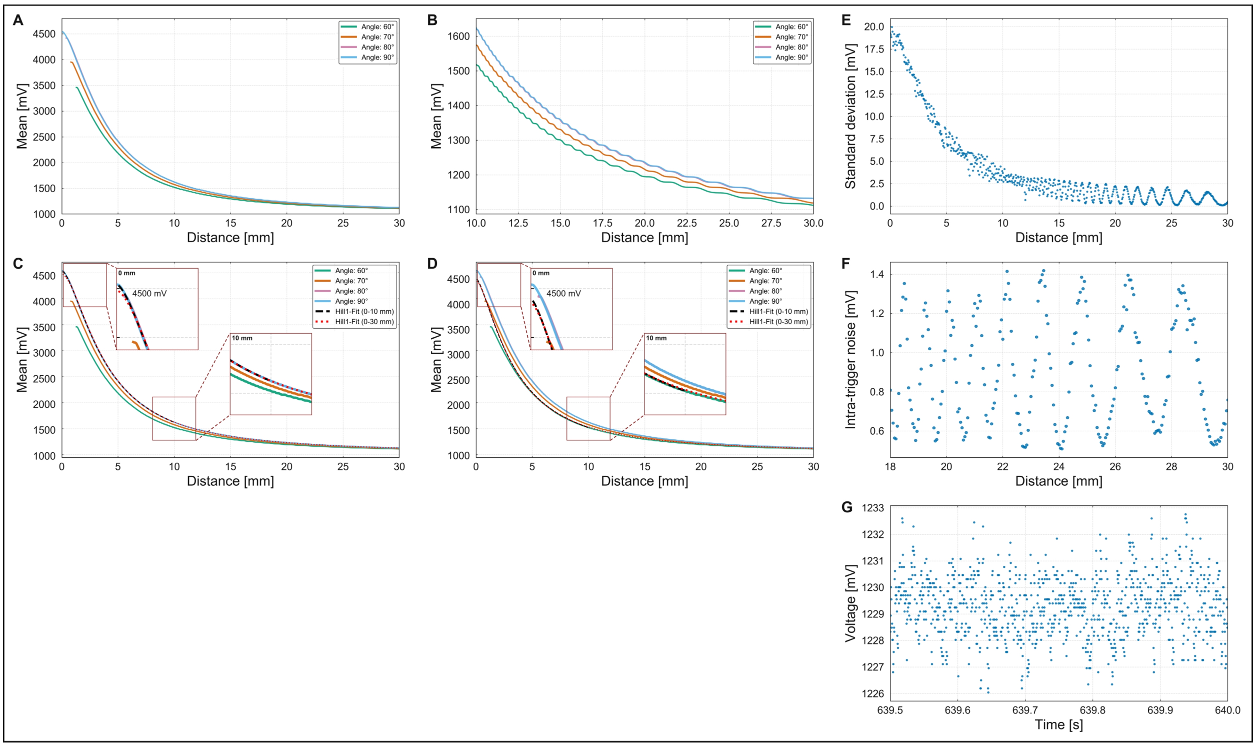


***Supplementary Figure 2:*** Angle-dependent voltage response, Hill1 calibration, and signal variability for Sensor 3. Voltage, distance, and time are reported in millivolts (mV), millimetres (mm), and seconds (s), respectively. **(A)** Mean monitor-output voltage across 30 repeated scans over 0–30 mm at incidence angles of 60°, 70°, 80°, and 90°. **(B)** Enlarged view of the 10–30 mm range, showing the distance-dependent modulation at larger separations. **(C)** Mean voltage–distance responses with uncertainty-weighted Hill1 fits derived from the pooled 80° and 90° data. **(D)** Mean responses with uncertainty-weighted Hill1 fits derived from the pooled 60°–90° data. Black dashed lines represent fits over 0–10 mm, whereas red dotted lines represent fits over 0–30 mm. In panels C and D, the dark-red boxes mark the selected regions, the dashed connectors link these regions to the corresponding insets, and the upper and lower insets enlarge the regions around the nominal 0 and 10 mm positions, respectively. **(E)** Standard deviation of the mean voltage across 30 repeated scans at each distance over 0–30 mm at 90°. **(F)** Intra-trigger noise, calculated as the standard deviation of approximately 40 samples within each 20 ms trigger window and averaged across the 30 scans, shown over 18–30 mm at 90°. **(G)** A 0.5 s segment of the monitor-output voltage at 20 mm and 90°, showing the sampled voltage levels over time. The measured 50° data are not displayed because these measurements were excluded from the pooled calibration fits according to the criterion described in the Methods.

Supplementary Figure 3 — Sensor 4

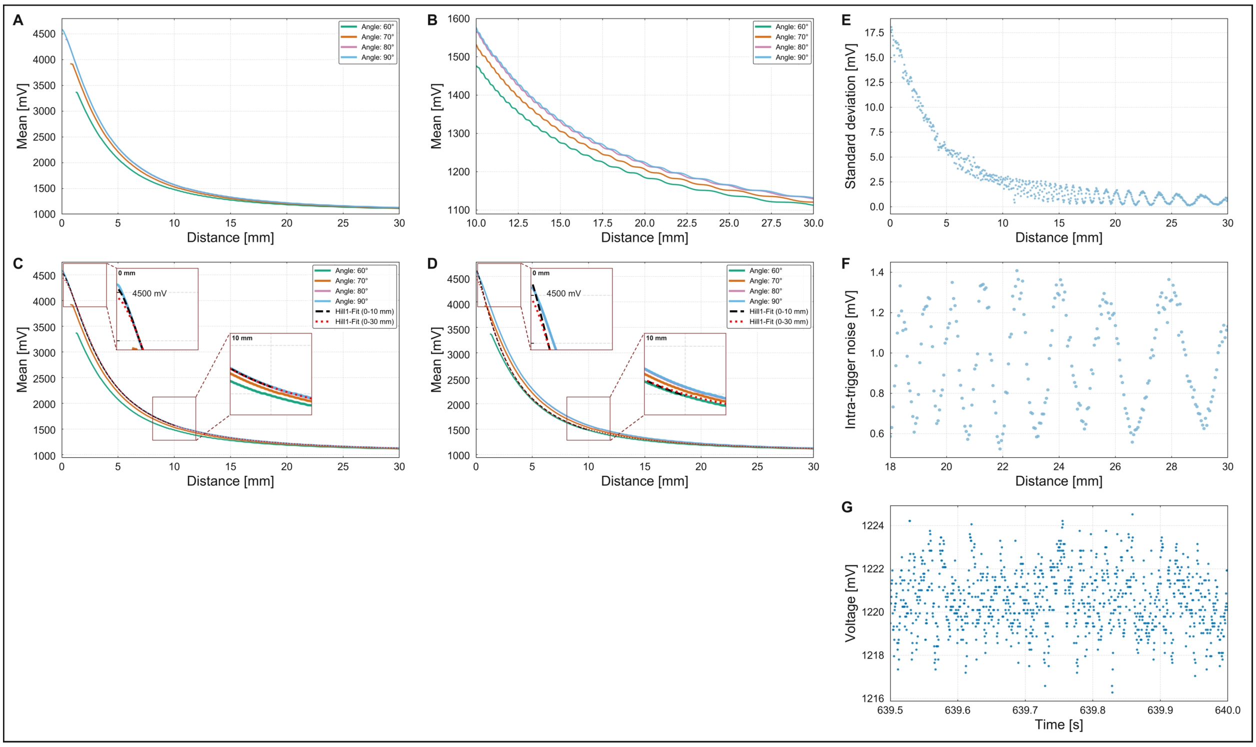


***Supplementary Figure 3:*** Angle-dependent voltage response, Hill1 calibration, and signal variability for Sensor 4. Voltage, distance, and time are reported in millivolts (mV), millimetres (mm), and seconds (s), respectively. **(A)** Mean monitor-output voltage across 30 repeated scans over 0–30 mm at incidence angles of 60°, 70°, 80°, and 90°. **(B)** Enlarged view of the 10–30 mm range, showing the distance-dependent modulation at larger separations. **(C)** Mean voltage–distance responses with uncertainty-weighted Hill1 fits derived from the pooled 80° and 90° data. **(D)** Mean responses with uncertainty-weighted Hill1 fits derived from the pooled 60°–90° data. Black dashed lines represent fits over 0–10 mm, whereas red dotted lines represent fits over 0–30 mm. In panels C and D, the dark-red boxes mark the selected regions, the dashed connectors link these regions to the corresponding insets, and the upper and lower insets enlarge the regions around the nominal 0 and 10 mm positions, respectively. **(E)** Standard deviation of the mean voltage across 30 repeated scans at each distance over 0–30 mm at 90°. **(F)** Intra-trigger noise, calculated as the standard deviation of approximately 40 samples within each 20 ms trigger window and averaged across the 30 scans, shown over 18–30 mm at 90°. **(G)** A 0.5 s segment of the monitor-output voltage at 20 mm and 90°, showing the sampled voltage levels over time. The measured 50° data are not displayed because these measurements were excluded from the pooled calibration fits according to the criterion described in the Methods.

### Supplementary Figure 4 — Sensor 5

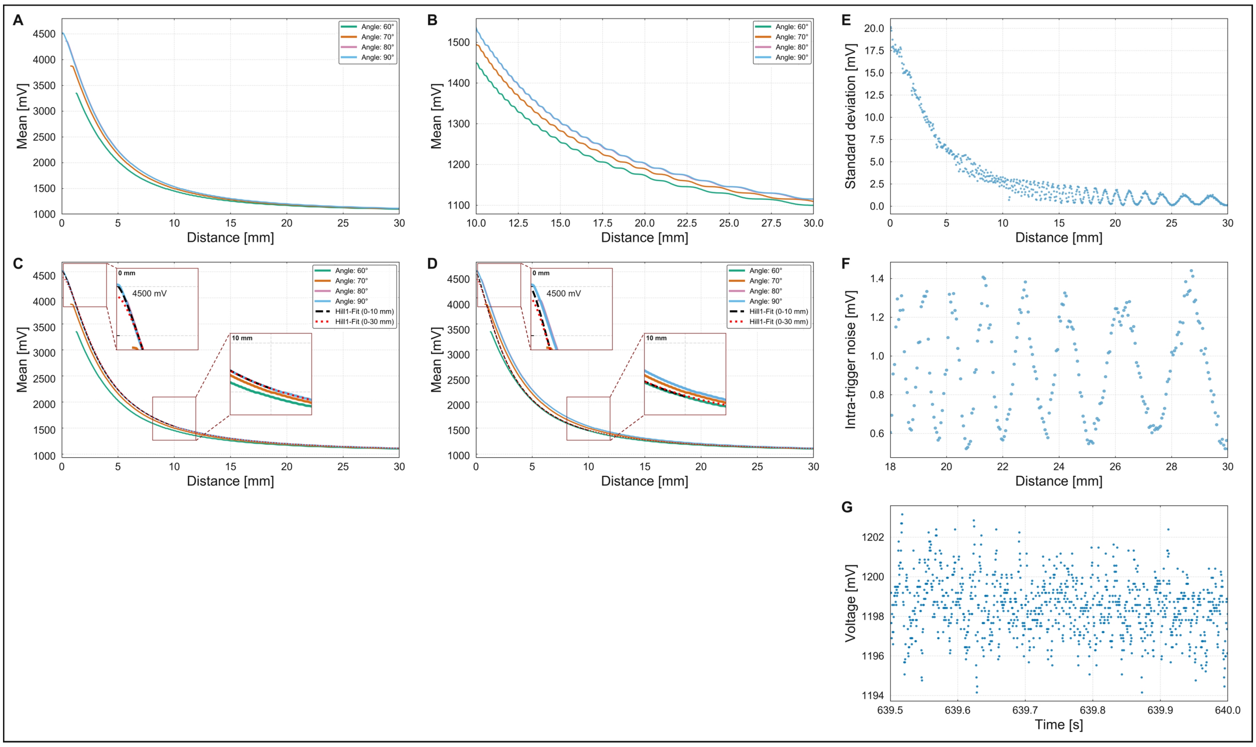


***Supplementary Figure 4:*** Angle-dependent voltage response, Hill1 calibration, and signal variability for Sensor 5. Voltage, distance, and time are reported in millivolts (mV), millimetres (mm), and seconds (s), respectively. **(A)** Mean monitor-output voltage across 30 repeated scans over 0–30 mm at incidence angles of 60°, 70°, 80°, and 90°. **(B)** Enlarged view of the 10–30 mm range, showing the distance-dependent modulation at larger separations. **(C)** Mean voltage–distance responses with uncertainty-weighted Hill1 fits derived from the pooled 80° and 90° data. **(D)** Mean responses with uncertainty-weighted Hill1 fits derived from the pooled 60°–90° data. Black dashed lines represent fits over 0–10 mm, whereas red dotted lines represent fits over 0–30 mm. In panels C and D, the dark-red boxes mark the selected regions, the dashed connectors link these regions to the corresponding insets, and the upper and lower insets enlarge the regions around the nominal 0 and 10 mm positions, respectively. **(E)** Standard deviation of the mean voltage across 30 repeated scans at each distance over 0–30 mm at 90°. **(F)** Intra-trigger noise, calculated as the standard deviation of approximately 40 samples within each 20 ms trigger window and averaged across the 30 scans, shown over 18–30 mm at 90°. **(G)** A 0.5 s segment of the monitor-output voltage at 20 mm and 90°, showing the sampled voltage levels over time. The measured 50° data are not displayed because these measurements were excluded from the pooled calibration fits according to the criterion described in the Methods.

**Supplementary Table S1.** Descriptive statistics for independent distance reconstruction errors across five sensors.

| Condition | Calibration range | Evaluation range | Error metric | n | Descriptive statistics across sensors (mm) | | | | |
|---|---|---|---|---|---|---|---|---|---|
| | | | | | **Mean** | **SD** | **Median** | **Minimum** | **Maximum** |
| 1 | 0–10 | 0–10 | Mean absolute error | 5 | 0.0282 | 0.0008 | 0.0280 | 0.0270 | 0.0290 |
| | | | Root mean squared error | 5 | 0.0358 | 0.0024 | 0.0360 | 0.0330 | 0.0390 |
| | | | Mean bias | 5 | -0.0182 | 0.0136 | -0.0230 | -0.0260 | 0.0060 |
| | | | Maximum absolute error | 5 | 0.1656 | 0.0560 | 0.1870 | 0.0860 | 0.2310 |
| 2 | 0–30 | 0–10 | Mean absolute error | 5 | 0.0396 | 0.0048 | 0.0390 | 0.0350 | 0.0470 |
| | | | Root mean squared error | 5 | 0.0528 | 0.0052 | 0.0540 | 0.0470 | 0.0590 |
| | | | Mean bias | 5 | -0.0140 | 0.0116 | -0.0200 | -0.0210 | 0.0060 |
| | | | Maximum absolute error | 5 | 0.3032 | 0.0716 | 0.3390 | 0.1880 | 0.3680 |
| 3 | 0–30 | 0–30 | Mean absolute error | 5 | 0.1010 | 0.0125 | 0.1000 | 0.0830 | 0.1170 |
| | | | Root mean squared error | 5 | 0.1446 | 0.0179 | 0.1410 | 0.1210 | 0.1700 |
| | | | Mean bias | 5 | -0.0460 | 0.0270 | -0.0540 | -0.0730 | -0.0010 |
| | | | Maximum absolute error | 5 | 0.6426 | 0.0745 | 0.6270 | 0.5840 | 0.7690 |
| 4 | 0–30 | 10–30 | Mean absolute error | 5 | 0.1310 | 0.0170 | 0.1290 | 0.1070 | 0.1540 |
| | | | Root mean squared error | 5 | 0.1724 | 0.0215 | 0.1680 | 0.1440 | 0.2030 |
| | | | Mean bias | 5 | -0.0616 | 0.0344 | -0.0700 | -0.0980 | -0.0050 |
| | | | Maximum absolute error | 5 | 0.6426 | 0.0745 | 0.6270 | 0.5840 | 0.7690 |

***Supplementary Table 1****:* The reconstruction analysis used the 80° and 90° datasets, which showed relatively high signal amplitudes and similar monotonic voltage–distance profiles in Fig. 2 and were therefore considered suitable for this initial evaluation. This selection was not intended to identify an optimal incidence angle. Scans 1–25 were used for Hill1 calibration, whereas scans 26–30 were retained as independent test data. Conditions 1–4 represent calibration/evaluation ranges of 0–10/0–10, 0–30/0–10, 0–30/0–30, and 0–30/10–30 mm, respectively. Reconstruction error was defined as the reconstructed distance minus the nominal distance. Mean absolute error (MAE), root mean squared error (RMSE), mean bias, and maximum absolute error were calculated for each sensor. Negative mean bias indicates a tendency to underestimate distance. The sample size n = 5 represents the five sensors. Mean, standard deviation (SD), median, minimum, and maximum were calculated across sensors. All distance and error values are reported in millimetres (mm).
